# Investigating the probability of establishment of Zika virus and detection through mosquito surveillance under different temperature conditions

**DOI:** 10.1101/406116

**Authors:** Rebecca C. Christofferson

## Abstract

Because of the increasing threat that Zika virus (ZIKV) poses to extra-tropical regions due to increased global travel, there is a need for better understanding of the effect(s) of temperature on the establishment potential of ZIKV within these subtropical, temperate, and/or seasonal Ae. aegypti populations. The first step to determining risk establishment of ZIKV in these regions is to assess ZIKV’s ability to infect mosquitoes at less tropical temperatures, and thus be detected through common surveillance programs. To that end, the effect of two rearing temperatures (RT) and extrinsic incubation temperatures (EIT) on infection and dissemination rates was evaluated, as well as the interactions of such. Total, there were four combinations (RT24-EIT24, RT24-DEI28, RT28-EIT24, RT28- EIT28). Further, a stochastic SEIR framework was adapted to determine whether observed data could lead to differential success of establishment of ZIKV in naive mosquito populations. There was no consistent pattern in significant differences found across treatments for either infection or dissemination rates (p>0.05), where only a significant difference was found in infection rates between RT24- EIT24 (44%) and RT28-EIT24(82.6%). Across all temperature conditions, the model predicted between a 77.3% and 93.1% chance of successful establishment of ZIKV in naive mosquito populations under model assumptions. Further, the model predicted between 4.1% and 46.7% chance of at least one mosquito developing a disseminated infection, depending on temperature conditions, despite no significant differences in the experimental data. These results indicate that 1) there is no straightforward relationship between RT, EIT, and infection/dissemination rates for ZIKV, similar to what has been reported for DENV, 2) in more temperate climates, ZIKV may still have the ability to establish in populations of Ae. aegypti, and 3) despite a lack of statistical differences in observed experimental data, model predictions indicate that the interplay of rearing and extrinsic incubation temperatures may still alter the kinetics of ZIKV within the mosquito enough to affect numbers of infected/disseminated mosquitoes and the associated probability of detection through surveillance programs.

## Introduction

Zika virus (ZIKV) emerged as a public health emergency in 2016 after a steady migration westward [1, 2]. As ZIKV spread quickly across the tropics of the Western Hemisphere, the primary mosquito vector was demonstrated to likely be *Ae. aegypti* which has efficiently transmitted dengue virus (DENV) in the region for decades [3, 4]. The ecology of ZIKV is similar to that of DENV and chikungunya (CHIKV) owing to the shared primary vector and transmission of these three viruses overlapped in several instances [5-7].

Annually, thousands of travelers from around the world contract these arboviruses, and a subset of these travelers return from areas of intense transmission to places with the potential for these viruses to establish in local mosquito populations [8-13]. A major reason these viruses have not emerged with the same intensity in extra-tropical regions is because of the distribution of *Ae. aegypti*, which is constrained by its affinity for tropical and sub-tropical climate [12]. However, there have been instances of seasonal transmission of DENV, CHIKV, and ZIKV in sub-tropical and even temperate areas [14-16]. Additionally, the CDC recently updated its predicted range for the principle ZIKV vectors, which shows a significant portion of the Eastern United States is at risk for at least a “likely” presence of *Ae. aegypti* [17].

This estimated distribution shows that the mosquito is likely to have at least some presence in more temperate regions, where there is a probability that development and potential viral incubation temperatures would be in the ranges of 24°C and 28°C. Indeed, a recent study found that *Ae. aegypti* were able to become infected at moderate rates at both 18°C and 27°C by 14 days post infection (dpi) suggesting there is not a definite temperature-driven barrier to the establishment of ZIKV in affected mosquito populations [18].

A better understanding of the ability and drivers of ZIKV to establish in mosquito populations that experience more temperate temperature profiles is needed to assess risk in these areas. It has been demonstrated that rearing temperatures can affect the infection and dissemination of arboviruses through *Ae. aegypti* [19-21]. However, the results are not uniform and indicate that these effects still need characterization [22, 23]. On the other hand, higher temperatures during the extrinsic incubation period (EIP) are almost uniformly associated with higher rates of infection and dissemination in many arboviral systems [18, 24, 25].

This study focused on the interplay of rearing temperature (RT) and extrinsic incubation temperature (EIT) at lower temperatures (24°C and 28°C) to determine whether there was an effect on the ability of ZIKV 1) to establish in mosquito populations and 2) to escape the midgut barrier and develop into a disseminated infection. These are two critical first steps in establishment of successful transmission chains. Further, whole-body mosquito pool testing is the most common method of detecting arboviruses in resident mosquito populations under usual surveillance programs. Therefore, a positive pool, which is the first indication of local Zika activity, is based on infected status only. Thus, a mathematical model was employed to determine the differential probabilities of establishment of infection in naïve mosquito populations – and thus detection –based on temperature treatments; and further the model predicts the number of disseminated infections from these results.

## Materials and Methods

### Mosquitoes and virus

Mosquitoes (Rockefeller strain) were originally obtained from Dr. Daniel Swale of the LSU Entomology department. The PRVABC59 strain of ZIKV was used and was originally obtained from the CDC originally isolated from a patient in Puerto Rico [26]. Viral titers of fresh (not frozen) ZIKV were confirmed by qRT-PCR and matched (~8 × 10^7^ PFU/mL) across same-day oral feedings using whole bovine blood with Alsevers (Hemostat Laboratories, Dixon, CA) and the Hemotek membrane feeding apparatus (Discovery Workshops, UK) as in [27, 28].

### Rearing and oral exposure of mosquitoes

Mosquitoes were allowed to hatch at one of two temperatures (24°C or 28°C) and kept at this temperature until they were exposed to an infectious blood meal at 3-5 days post emergence. This is hereafter referred to as the rearing temperature (RT). After exposure to ZIKV, fully engorged females were sorted into clean cartons. Half of these were left at their original RT for the extrinsic incubation period while the other half were moved to the opposite temperature. This temperature is the extrinsic incubation temperature (EIT). Thus, there were a total of 4 treatments: RT28-EIT28, RT28-EIT24, RT24-EIT28, and RT24-EIT24 (Figure S1).

### Sampling, processing, and detection of ZIKV from mosquitoes

To quantify the rates of infection and dissemination over time, mosquitoes were sampled on days 7, 10, and 13 post-exposure and tested for the presence of virus in the abdomens and legs, respectively. These time points were chosen based on studies that have demonstrated the timing of ZIKV within *Ae. aegypti* mosquitoes [29, 30]. Briefly, mosquitoes were cold anesthetized and the legs separated from the bodies and placed into 2mL tubes with 900 µl of BA1 and two steel-coated BBs. Samples were then homogenized using a TissueLyzer (Qiagen), centrifuged at 4,000 rpm for 4 minutes, frozen at −80°C. Samples were thawed and RNA extracted using the KingFisher (ThermoFisher) robot and the Ambion MagMax viral isolation kit (ThermoFisher), as per manufactuer’s instructions. Samples were tested for the presence of ZIKV RNA via qRT-PCR using the SuperScript III One-Step RT-PCR System with Platinum Taq DNA Polymerase (Life Technologies) on the Roche LightCycler 480 system using primers and probes previously published [31, 32]. Samples with a CT of <35 were considered positive, while samples with a CT of 35 or greater were inoculated onto Vero cells and the supernatant tested 4 days later. At this point, CT<35 was indicative of a positive sample, but a second CT value of 35 or higher was indicative of a negative sample.

### Statistical analysis of experimental data

Data were analyzed using RStudio (version 1.1.383, with base R version 3.4.3). Mosquito infection and dissemination rates were compared daily by chisquare test for equivalency of frequencies using the R command prop.test (with continuity correction, [33]). Infection rates were calculated as the number with a positive body divided by total number exposed. Dissemination rates were first calculated as the number with positive legs divided by the total number tested for a measure of population-level dissemination, and then as the proportion of disseminated infections out of total positive bodies (infected). Presence of virus in the legs indicates that the virus has gotten out of the midgut, the first within-mosquito barrier to transmission.

### Modeling the temporal processes of infection and dissemination

To further describe these processes, the data was fit using a non-linear least squares model, to determine the best exponential fit (R version 3.4.3) as described in [34]. An origin at (0,0) was assumed and to satisfy non-zero requirements of data, 1×10^-6^ was added to all data points. Briefly,

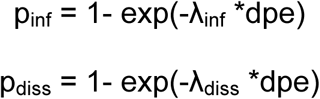

where p_inf_ is the proportion of samples with positive bodies, λ_inf_ is the rate parameter of the exponential cumulative distribution function (CDF); p_diss_ is the proportion of mosquitoes that developed a disseminated infection given they were infected (disseminated/infected); λ_diss_ is the rate parameter fit to the disseminated/infected data; and dpe is the day post exposure corresponding to each value p [34]. Following estimates, the fit was assessed via Kolmogorov– Smirnov test for goodness of fit by comparing a randomly generated exponential CDF with the parameter estimates from the experimental data to the data itself and all estimated fits were deemed “good” as p>0.05, indicating no rejection of the null hypothesis that no differences exist between the two distributions.

Subsequently, a stochastic, SEIR compartmental model was modified from [35]. Briefly, the model incorporated the values of λ_inf_ to describe the average rate of movement of mosquitoes from the exposed to the infected class, and λ_diss_ describes the rate of movement from infected to the disseminated class [34]. These rates of movement were further weighted by the maximum proportion of infected or disseminated mosquitoes so that exposed →infected was defined as p_inf.max_*λ_inf_ and infected → disseminated was defined as p_diss.max_*λ_diss_.

Because the purpose of this exercise was to describe the differential potential of ZIKV to establish in mosquitoes at different temperature profiles, and the probability of detection in mosquito pools (i.e. a single infected mosquito), the model did not allow mosquitoes to transmit, and the model was run for a total of 30 days (Figure 1). That is, the final output is number of mosquitoes exposed, infected, and disseminated following a primary, single introduction of five infectious humans (returning travelers, e.g.) and accounts for a single generation of infection (human → mosquito). Subsequently, the probability of ZIKV establishing infection in or disseminating through at least one mosquito was calculated as the number of simulations with at least one infected or disseminated mosquito (respectively) divided by the total number of simulations. Model details are given in the Supplemental Information, Supplemental Table 1, and Supplemental Figure 1. A total of 500 simulations per treatment were run.

**Table 1:** Parameter values (rounded to nearest hundredth) determined by non-linear least squares (λ) and the maximum proportion (p_max_) of infection or dissemination determined by the experimental data.

| Treatment | $\lambda$ | | $p_{\max}$ | |
| --- | --- | --- | --- | --- |
| RT24-EIT24 | $\lambda_{\text{inf}}$ | 1/14.01 | $p_{\text{inf}}$ | 0.58 |
| | $\lambda_{\text{diss}}$ | 1/70 | $p_{\text{diss}}$ | 0.21 |
| RT24-EIT28 | $\lambda_{\text{inf}}$ | 1/11.7 | $p_{\text{inf}}$ | 0.71 |
| | $\lambda_{\text{diss}}$ | 1/14.6 | $p_{\text{diss}}$ | 0.67 |
| RT28-EIT24 | $\lambda_{\text{inf}}$ | 1/6.3 | $p_{\text{inf}}$ | 0.89 |
| | $\lambda_{\text{diss}}$ | 1/43.3 | $p_{\text{diss}}$ | 0.28 |
| RT28-EIT28 | $\lambda_{\text{inf}}$ | 1/8.5 | $p_{\text{inf}}$ | 0.79 |
| | $\lambda_{\text{diss}}$ | 1/17.9 | $p_{\text{diss}}$ | 0.64 |

**Figure 1:**
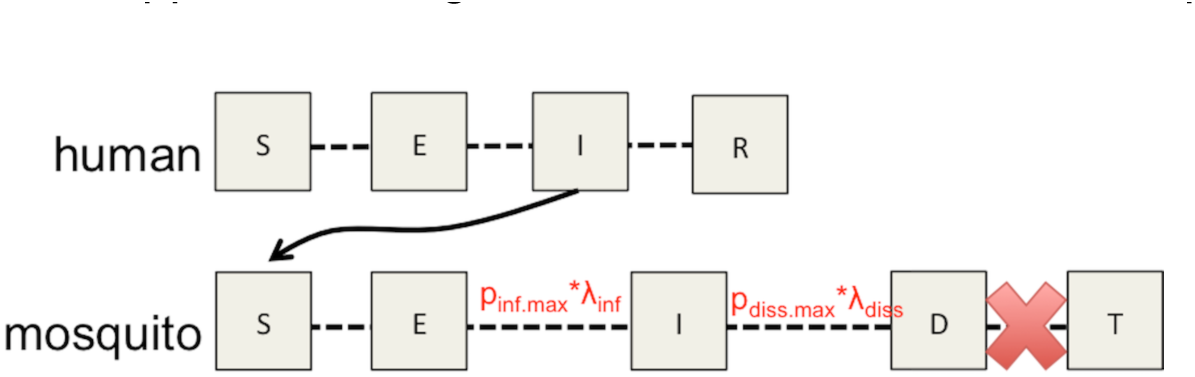
Model schematic demonstrating how experimental data informs the parameterization of a model to simulate the probability of ZIKV establishing infection in at least one mosquito and predicting the probability of at least one disseminated infection following introduction into a naïve population of *Ae. aegypti*.

## Results

### Effect of RT-EIT on infection and dissemination rates

The infection and dissemination rates are shown in Figure 3. At days 7- and 13-post infection, there was no significant difference in either the ability of ZIKV to infect or to disseminate through mosquitoes among temperature combinations (p>.05). At day 10, there was a significant difference in the proportion of infected mosquitoes, and pairwise comparisons identified this difference between RT24-EIT24 (44%) and RT28-EIT24 (82.6%). However, there was observed no significant difference in the ability of ZIKV to escape the midgut among the temperature combinations. When the proportion of disseminated mosquitoes was calculated as the proportion positive divided by infected mosquitoes, there was a significant difference at day 7 and day 13. However, when the proportion of disseminated mosquitoes was calculated as the proportion positive divided by infected mosquitoes, there was a significant difference at day 7 and day 13. At 7 dpe, there was a significant difference between RT24-EIT24 and RT24-EIT28 and between RT24-EIT28 and RT28- EIT24, where the higher EIT resulted in higher dissemination. Subsequent pairwise comparisons at day 13 did not result in any statistical significant (Figure 2).

**Figure 2:**
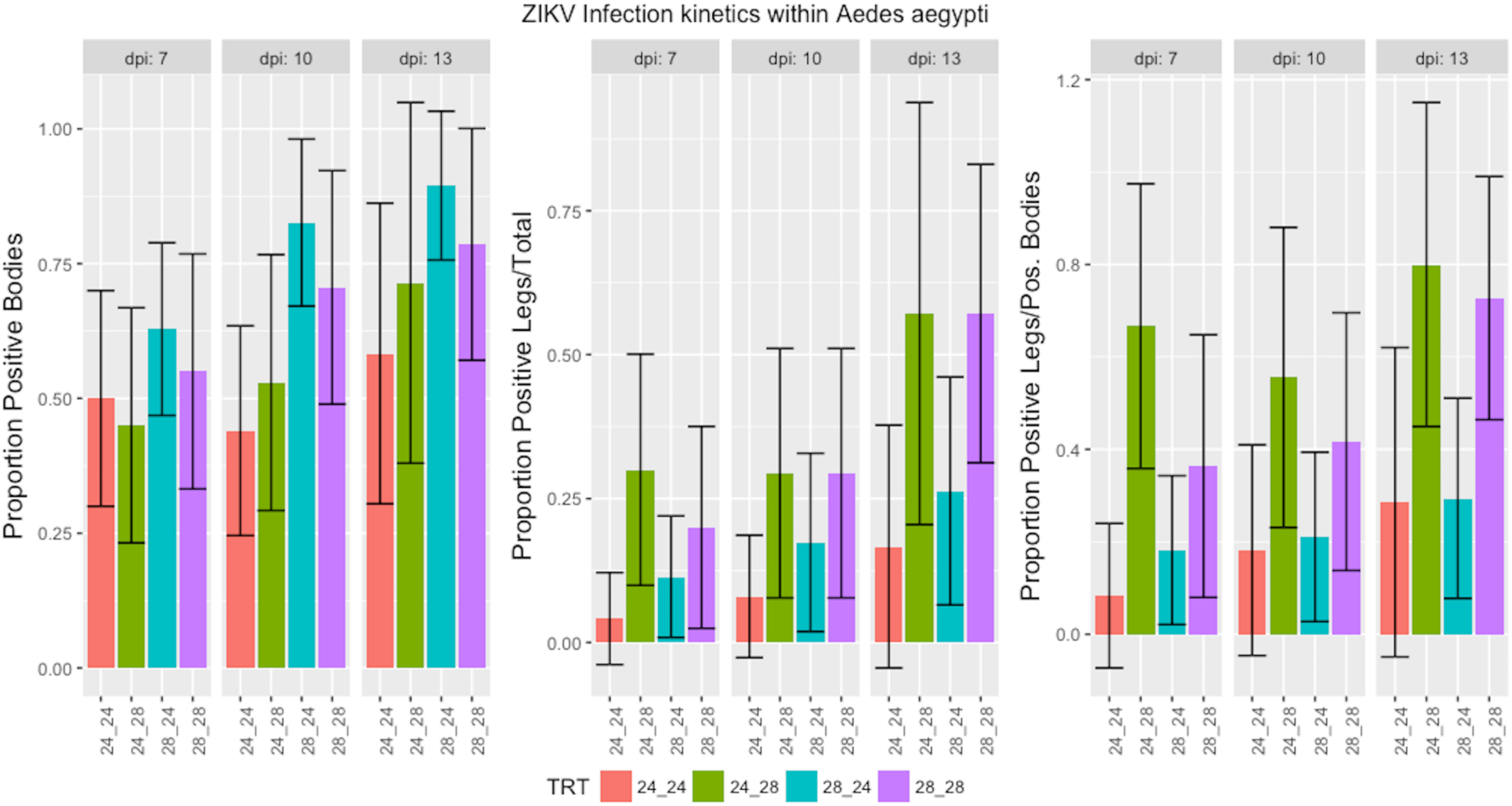
Proportion of mosquitoes that became (Left) infected with Zika, and those that developed a disseminated infection calculated as (Middle) disseminated/total and (Right) disseminated/infected. Statistical significance was determined via Chi-square test for equal proportions (α=0.05). Error bars represent the binomial 95% confidence intervals of the proportions.

**Figure 3:**
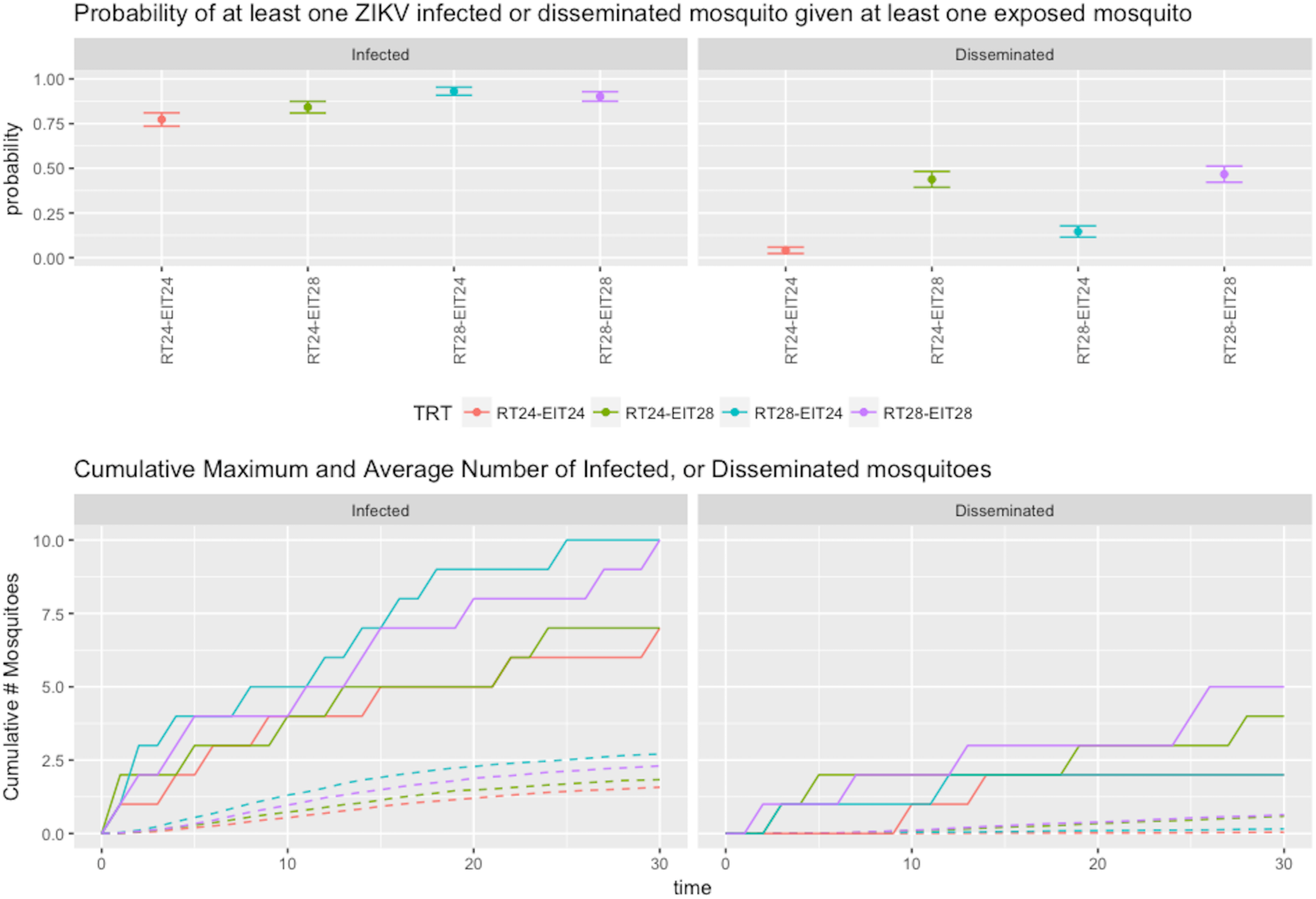
**Top** – The probabilities of at least one infected (left) or disseminated (right) mosquito given at least one exposed mosquito following introduction of ZIKV infected humans into a naïve mosquito population.**Bottom** – The simulated cumulative maximum (solid lines) and average cumulative (dotted lines) number of infected and disseminated mosquitoes over the course of 30 days following the introduction of 5 infectious humans into a naïve mosquito population.

### Parameterization of compartmental model

Table 1 shows the values of λ determined through non-linear least squares modeling using nls function in R and the maximum values for infection and disseminated (out of total infected) proportions from the experimental data. In the model, the transmission rate was set at 0 as the purpose was to identify establishment after a single generation of human-to-mosquito transmission (and not mosquito-to-human back to mosquito). There were no additional human infections other than the initial value of five. Thus the model is appropriate to assess mosquito-only infection kinetics.

For each set of temperature conditions, there was between a 95.6% and 96.8% probability of at least one mosquito becoming exposed under model conditions and given an initial introduction of five infected humans (Figure S1). Thus, the number of simulations that failed to produce any exposed mosquitoes was less than 5% for all treatment conditions. When examining the binomial confidence intervals of these probabilities, there were no differences among predicted probabilities of at least one exposed mosquito, indicating the model is appropriate to evaluate post-exposure kinetics.

To assess ZIKV kinetics of infection and dissemination within the population, probabilities were calculated as the number of simulations where at least one mosquito became infected/disseminated divided by the total number of simulations where at least one mosquito had become exposed (as described above). There was between a 77.3% and 93.1% chance that ZIKV would successfully establish infections in mosquitoes (Figure 3). The lowest probability of infection was not surprisingly in the RT24-EIT24 group while the highest chance of successful establishment was in the RT28-EIT24 group. However, the overlap of confidence intervals of RT28-EIT24 and RT28-EIT28 (90.2%) likely means this difference is due to stochastic processes. The RT24-EIT28 group had a probability of established infection of 84.2%. Overall, there was overlap of the 95% binomial confidence intervals among most treatments, signifying likely no significant pattern of infection success and temperature conditions.

When dissemination was predicted given at least one exposed mosquito, the highest probabilities were in the RT28-EIT28 group (46.7%) and the RT24- EIT28 group (43.8%). There was a moderate probability of dissemination in the RT28-EIT24 group (14.6%) and a low probability in the RT24-EIT24 group (4.1%). These results indicate that higher EITs are more important predictors of successful dissemination.

The maximum number of mosquitoes at each time point was summed cumulatively over the 30 days modeled for each set of temperature conditions. Simulations predicted that the cumulative maximum number of infected mosquitoes ranged from 7 (RT24-EIT24, RT24-EIT28) to 10 (RT28-EIT24, RT28-EIT28) (Figure 3). The number of mosquitoes predicted to develop a disseminated infection was 2 (RT24-EIT24, RT28-EIT24), 4 (RT24-EIT28), and 5 (RT28-EIT28). The average cumulative number of infected and disseminated mosquitoes is given in Figure 3 and was calculated as the average out of the total simulations resulting in at least one exposed mosquito.

## Discussion

After a mosquito takes a bloodmeal, any arbovirus in that bloodmeal must establish infection in the midgut. The virus must then get past the midgut barrier in order to disseminate to the peripheral tissues. These are both critical first steps in the process of establishing a chain of transmission in naïve populations, as failure to infect and subsequently get past the midgut barrier means that mosquito will not be able to transmit. Temperature is a known driver of vector competence, but whether it acts directly on this midgut barrier is unknown though data suggests higher EITs lead to faster and higher rates of dissemination [18, 24, 25]. Less characterized is the role of larval temperature conditions on subsequent vector competence. While there is some lack of agreement among the effects of RT and EIT on the reported experimental ZIKV kinetics and the simulated infection probabilities across the four treatments, the prediction of dissemination success follows the tenant that higher EITs leads to more disseminated infections and could potentially validate higher transmission rates. This lack of a consistent relationship between temperatures experienced during the juvenile and adult stages has been observed in other arbovirus systems.

In one study of DENV-1, it was shown that both EIT and RT affected infection and dissemination rates at 14 dpe, but in a non-uniform manner [23]. However, in another study, rearing temperature (24°C vs 28°C) did not significantly affect DENV-1 infection rates of *Ae. aegypti* at 17 dpe [22]. The results of this study show that at 24°C, there is not a difference in the within-*aegypti* kinetics of ZIKV compared to 28°C, which is near-to the gold standard for vector competence studies [28, 29, 36-38]. While not statistically significant, the results of this study would seem to confirm the positive association of higher temperatures – especially during the EIP – with a trend towards higher and faster infection and dissemination rates [18, 24, 39]. But, importantly, it was demonstrated that ZIKV can infect and disseminate through mosquitoes at lower temperatures, and that fluctuations in temperature between juvenile and adults stages does not appear to alter vector competence.

Understanding the temporal nature of these processes compliments reporting of the magnitude of differences and can offer additional insights into the transmission dynamics of these viruses and is a useful and complimentary metric in describing within-host viral kinetics that is directly translatable to predictive models for sub-tropical and temperate regions of the world [34, 40-42]. The incorporation of this experimental data into the mathematical model was straightforward and appears to capture the relationships seen among the vector competence data reported and the transmission patterns expected from those data. Future studies are needed to assess the interactions of larval and adult habitat conditions on the propagation of ZIKV through multiple transmission generations, including actual transmission rates and associated life-traits that may or may not be temperature and/or infection dependent [28].

While use of the exponential fit to the data allows for direct translation of the data to the mathematical model, other transmission systems may be better fit by other distributions. Sometimes the data follow a sigmoid curve and in this case, a Gamma distribution would 1) fit the data appropriately, and 2) be relatively straight-forward to translate into a compartmental model such as the one developed for this study. However, comparisons of statistical distributions,data fitting, and mathematical model parameter estimation is beyond the scope of this study.

In conclusion, these results indicate that 1) there is no straightforward relationship between RT, EIT, and infection/dissemination rates, 2) in more temperature climates, ZIKV may still have the ability to establish in populations of *Aedes aegypti*, and 3) despite an overall lack of significant differences in infection/dissemination rates, temperature may still alter the kinetics of ZIKV within the mosquito enough to affect predicted numbers of infected/disseminated mosquitoes and the likelihood of detection within the context of mosquito surveillance programs. This suggests that, even though the cumulative maximum number of mosquitoes expected to be infected and/or disseminated in the first generation of transmission is low, there is some chance of successful transmission chains to be established.

### Declarations

*Ethics approval and consent to participate:* Not applicable

*Consent for publication:* Not applicable

## Acknowledgements

Thank you to Dr. Daniel Swale for provision of mosquito eggs and the Dr. Barbara Johnson at the CDC for Zika virus strain PRVABC59. Thank you to E. Handly Mayton and Kelly Hughes for their technical assistance and for being all-around good sports about most things, except spiders. Thank you to Dr. Michael McCracken for his valuable comments during the writing of this manuscript, and to Dr. Helen Wearing for her invaluable feedback, as per usual. Also, sorry to the spiders.

---

**Supplemental Figure 1**: Experimental Setup: Mosquito were reared at one constant temperature (either 24°C or 28°C). “Rearing Period” includes egg hatching, larval, pupal, and very young adult stages (up to 5 days post emergence). “Extrinsic Incubation Period (EIP)” refers to the time following ZIKV-infected blood-meal until sampling; the temperature associated with EIP (either 24°C or 28°C) is called the extrinsic incubation temperature (EIT).

**Supplemental Figure 2**: Probabilities of at least one exposed mosquito given the introduction of ZIKV infected humans into a naïve population.

